# Feasibility of adopting water displacement and 3D scanning methods to quantify lung volume reduction in *ex vivo* models

**DOI:** 10.64898/2026.09.02.748797

**Authors:** Ayla N. Kwant, Stacy L.S. Yam, Jan Aart M. Schipper, Maaike van Dillewijn, Adrivit Mukherjee, Joep Kraeima, Dirk-Jan Slebos, Janette K. Burgess, Simon D. Pouwels

## Abstract

Lung volume reduction is a treatment for chronic obstructive pulmonary disease (COPD) patients with severe emphysema. Currently, *in vivo* animal models are used as a first step to quantify the efficiency of novel lung volume reduction treatments. By quantifying lung volume reduction in *ex vivo* samples, animal testing may be reduced. This study proposes two new methods for quantifying lung volume reduction in *ex vivo* tissue: water displacement and three-dimensional (3D) scanning. Porcine lung lobes were inflated and treated with cyanoacrylate glue to simulate lung volume reduction. Volume changes were measured using a custom water displacement setup and in a ventilated setting using a handheld Artec Space Spider 3D surface scanner. Both techniques detected post-treatment volume reductions. Water displacement offers a simple, cost-effective approach for isolated lung lobes, while 3D scanning enables volumetric analysis of intact lungs under ventilated conditions. These methods provide a reliable intermediate step between *in vitro* and *in vivo* testing, reducing animal use and improving preclinical assessment of novel lung volume-altering interventions.

## Introduction

Assessing the volume of lung tissue in an *ex vivo* setting can be useful for pre-clinical studies. For instance, tissue volume measurements are useful for studying lung tumors, edema or fluid accumulation, or to assess the effectiveness of a lung volume-reduction treatment. To date, there are no studies that describe methods for accurately determining lung volume *ex vivo*. Only visual estimation has been used to describe the volume of lung tissue before and after a lung tissue volume reduction treatment [1]. Currently, the first step to study lung volume reduction is using *in vivo* animal such as pigs and sheep, measuring lung volume using CT after treatment with valves or adhesives[2,3]. In the current study, we describe two novel methods for quantifying lung volume in an *ex vivo* setting. Both methods can be used for a variety of situations, e.g. human lung tissue, which is available for *ex vivo* testing derived from tumor resection or lung volume reduction surgery or lung transplantation, whilst animal lung tissue is readily available as a waste product of the meat industry or animal experiments. Using these methods will provide opportunities to reduce animal usage through reuse of material that would traditionally be considered waste and to evaluate novel lung tissue reduction treatments more accurately.

Bronchoscopic lung volume reduction (BLVR) treatment is a minimally invasive therapeutic approach aimed at reducing lung volume in chronic obstructive pulmonary disease (COPD) patients with severe emphysema. By targeting hyperinflated regions of the lung, BLVR enables healthier lung tissue to expand and function more efficiently, thereby improving patient outcomes [4,5]. BLVR treatments include implanting endobronchial valves (EBVs), coils or the use of adhesives to seal emphysematous regions [6]. These interventions have proven effectiveness in enhancing lung capacity and improving the quality of life for COPD patients [4,5]. However, new BLVR treatments with improved efficacy and fewer adverse effects are still under development.

A key factor to study BLVR efficacy is to determine the volume of the targeted lung lobe before and after treatment. To avoid excessive animal testing, initial tests may be performed on *ex vivo* lung tissue. To accurately determine the volume reduction capacity of newly developed BLVR treatments, a reliable and quantitative method to determine *ex vivo* target lobe volume is required. Water displacement, based on the principle of Archimedes, is the gold standard for assessing volumes. It has been successfully applied to determine the volume of tumors and various soft tissues [7,8]. Moreover, water displacement has been used to study the volume of preserved lungs over time, offering a simple and cost-effective approach [9]. However, it only provides volume data of objects as a whole and does not allow specific or localized measurements. Recently, 3D scanning has emerged as an accurate new technique for determining the volume of breast tissue, tumors and limbs [8,10]. Hand-held, structured light-based scanners are particularly well-suited for laboratory use, and have been shown to reliably and accurately measure the volume of arms, legs and faces [11,12]. The Artec Space Spider is recommended for this application, as it has the smallest error margins (0.04 mm) when reliability was assessed [13]. To date, 3D scanning has never been utilized on *ex vivo* lung tissue. Both water displacement and 3D scanning may provide valuable insights into the efficacy of novel BLVR techniques. In this study, both methods will be used to quantify the volume of *ex vivo* lung tissue, focusing on the treated lobe. Water displacement will be used to determine the volume upon inflation of individual porcine lung lobes, before and after a volume reduction treatment with cyanoacrylate glue. Furthermore, BLVR treatments will be performed with intact *ex vivo* lungs, simulating a more clinically relevant scenario, in which the inflated, targeted areas will be 3D scanned before and after treatment. Our goal is to establish accurate and reproducible quantitative techniques for the early-stage evaluation of novel BLVR strategies that are more reliable than qualitative visual estimation alone.

## Methods

### Water displacement method

A glass beaker was glass-blown in-house (Figure 1A, diameter: 75mm; height: 100mm), with a spout (diameter: 8mm) at 15mm from the top to allow the displaced water to flow out, adapted from Rush *et. al*. [14]. A beaker was placed below the end of the spout to collect the displaced water, which was subsequently transferred to a graduated cylinder for volume measurement. Porcine lungs were obtained from a local slaughterhouse (Kroon Vlees, Groningen, The Netherlands). Lower lung lobes were secured into place by bamboo sticks that were placed into slits in the glassware to prevent the tissue from floating. The bottom of the slits were 5mm above the bottom of the spouts, keeping the amount of tissue above water constant. The transverse plane (Figure 1B) that exposes the airway was positioned facing upwards, and the remaining part of the lobe body was submerged in saline (0.9% NaCl in H_2_O).

**Figure 1.**
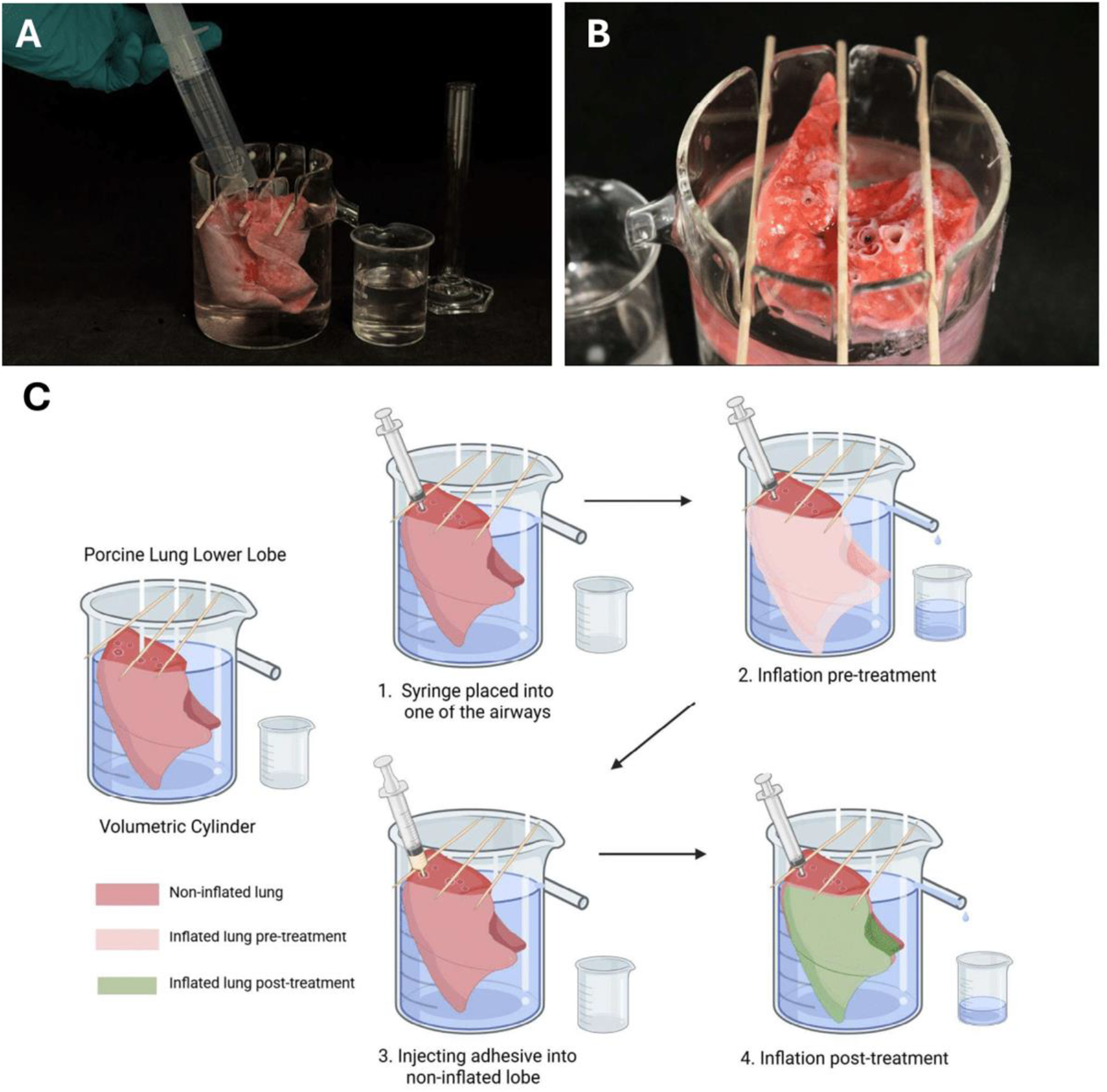
Water displacement method to measure ex vivo lung volume reduction. A) Experimental setup with a customized beaker with a lung lobe in place. B) Top view of the air-exposed plane of the porcine lung lobe with visible airways, allowing access for inflation and treatment using a syringe. C) Schematic representation of (partial) lung volume quantification method.

Before each measurement, the beaker was filled with saline exactly to the level of the spout of the beaker. The tip of a 20mL syringe was fitted tightly into the airway, and 20mL of air was injected to inflate the lobe, followed by measuring the displaced water volume. 0.5-1mL of cyanoacrylate glue was pipetted into the lobe via the same airway that was used for inflation. The amount of glue required was dependent on the anatomy of the lobe. Each administration filled up the tissue, while leaving room at the top of the airway for a syringe for later inflation. The glue was left to solidify for 3 minutes, after which the lobe was inflated again. All inflation and volumetric measurements were performed on 3 separate lung lobes with 5 technical replicates (5 volume measurements before treatment, then 5 volume measurements after treatment).

### 3D scanning method

The second volumetric measurement is based on a ventilated *ex vivo* porcine lung model and a handheld Artec Space Spider 3D scanner (Artec Europe Sarl, Luxembourg) (Figure 2). The porcine lung was intubated with an endotracheal tube and connected to a portable ventilator, Dräger Oxylog® 3000 plus (Dräger, Germany) and an Ambu® aScope ™ 4 Broncho bronchoscope (Ambu A/S, Denmark). The ventilator was set to volume control mode with a respiratory rate of 15 breaths per minute, positive end-expiratory pressure of 5 cm H_2_O, tidal volume of 750mL, maximum inspiratory pressure of 30 cm H_2_O, and oxygen concentration of 0.21 from the ambient air. Additionally, a vacuum was connected to the bronchoscope and was then used to visualize the airway environment and to clear airway secretions. The distal end of the bronchoscope contained a camera and a light source. After selecting a location in the distal airway for treatment, a Huibregtse ® 6-Fr guiding catheter (Cook Medical, United States) was inserted via the working channel of the bronchoscope to deliver the treatment. Sensitivity during scanning was set to maximum in the software (Artec Studio 14, Artec 3D, Luxembourg) to be able to capture the shape of the lung. The lung was scanned during its fully inflated state, while the ventilator was temporarily set to inspiratory hold mode. Inspiratory hold is important, as to not induce any movements while the scanning is performed, which may lead to inaccuracies in the results. The scanning was performed by slowly moving the scanning device within a distance of 0.2-0.3 meters, focusing on the treatment target area and surrounding areas. Both treated and non-treated tissue were scanned to facilitate registration of surfaces by matching the non-treated tissue. The ventilation was paused to deliver 1mL of cyanoacrylate glue through a catheter inserted through the bronchoscope working channel, which was left to solidify for 3 minutes. The ventilation was restarted with cyclic ventilation first to confirm the reduced recruitment to the treated region, followed by inspiratory hold mode, for 3D scanning of the treated lobe. It is important to note that the lungs were placed on a flat table during the entire experiment, meaning the bottom side is not scanned.

**Figure 2.**
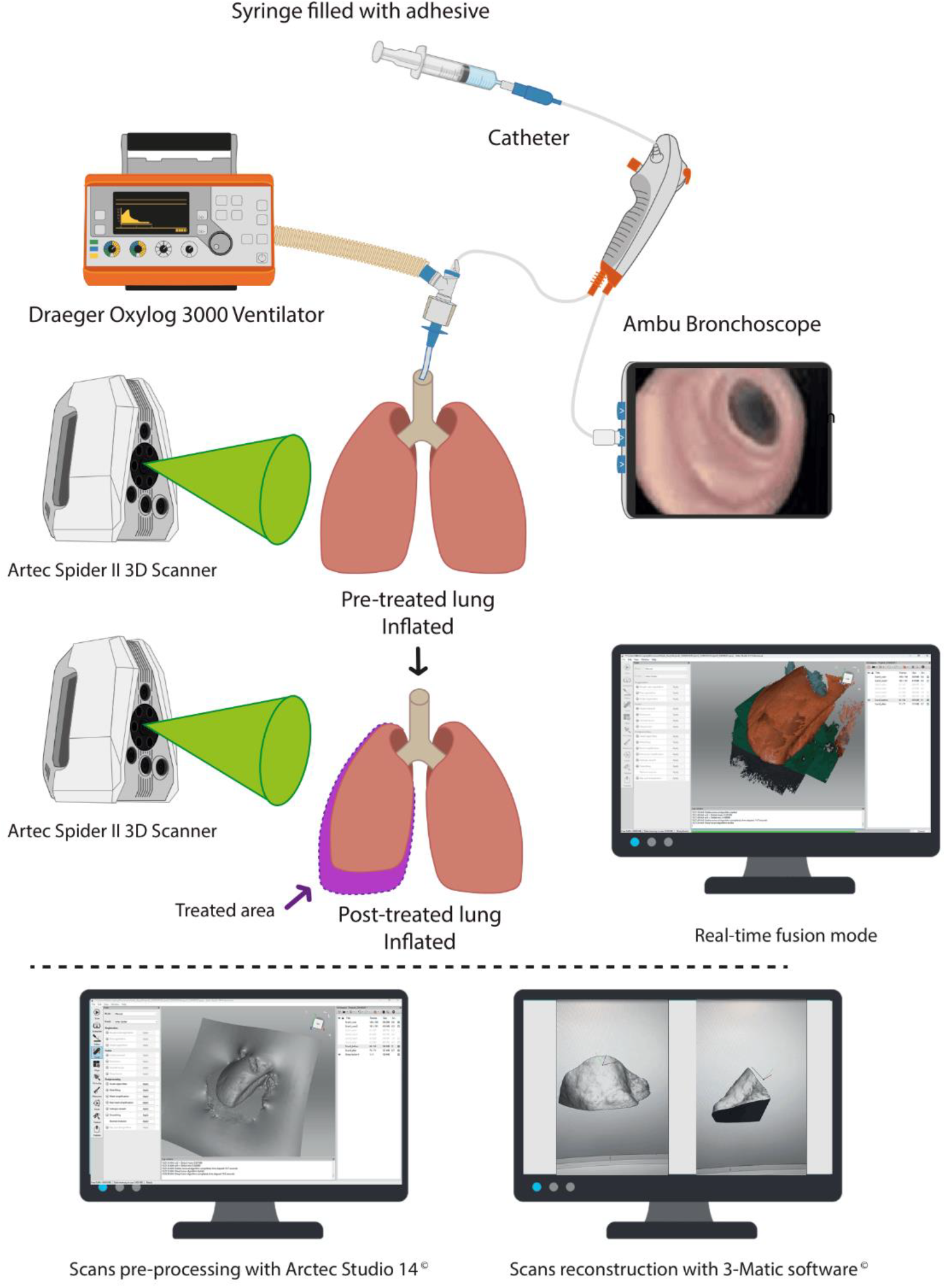
3D scan reconstruction method to measure lung volume reduction in the treated region. The lungs are inflated using the ventilator, and 3D scanned before treatment. Then, treatment is administered through the bronchoscope catheter, after which the same area of the lung is 3D scanned again. Processed and reconstructed scans are used to calculate the volume of the target area before and after treatment.

After capturing, the scans were first pre-processed using the Artec Studio 14 software. Scans were rendered into a 3D model by following standardized processing steps based on an earlier study [13]. Scans were then further analyzed with 3-matic software (Materialise, Leuven, Belgium). Registration *i*.*e*. alignment of the pre- and post-treatment scans was performed by matching the untreated regions that maintained a consistent shape during the procedure. After alignment, the pre- and post-treatment 3D surfaces were cropped, to focus on the targeted area. After cutting and removing all irrelevant areas, the surfaces were reconstructed into solid objects, and the volume difference was calculated automatically in the 3-matic software.

## Results

The volume of three porcine lung lobes was measured using the water displacement method before and after treatment (Figure 3A). Despite each lobe being inflated using 20mL of air, the measured inflated volume before treatment differs between lobes due to biological variation. All three lobes displayed a significant reduction in volume post-cyanoacrylate glue treatment (p=0.0079 for all lobes).

**Figure 3.**
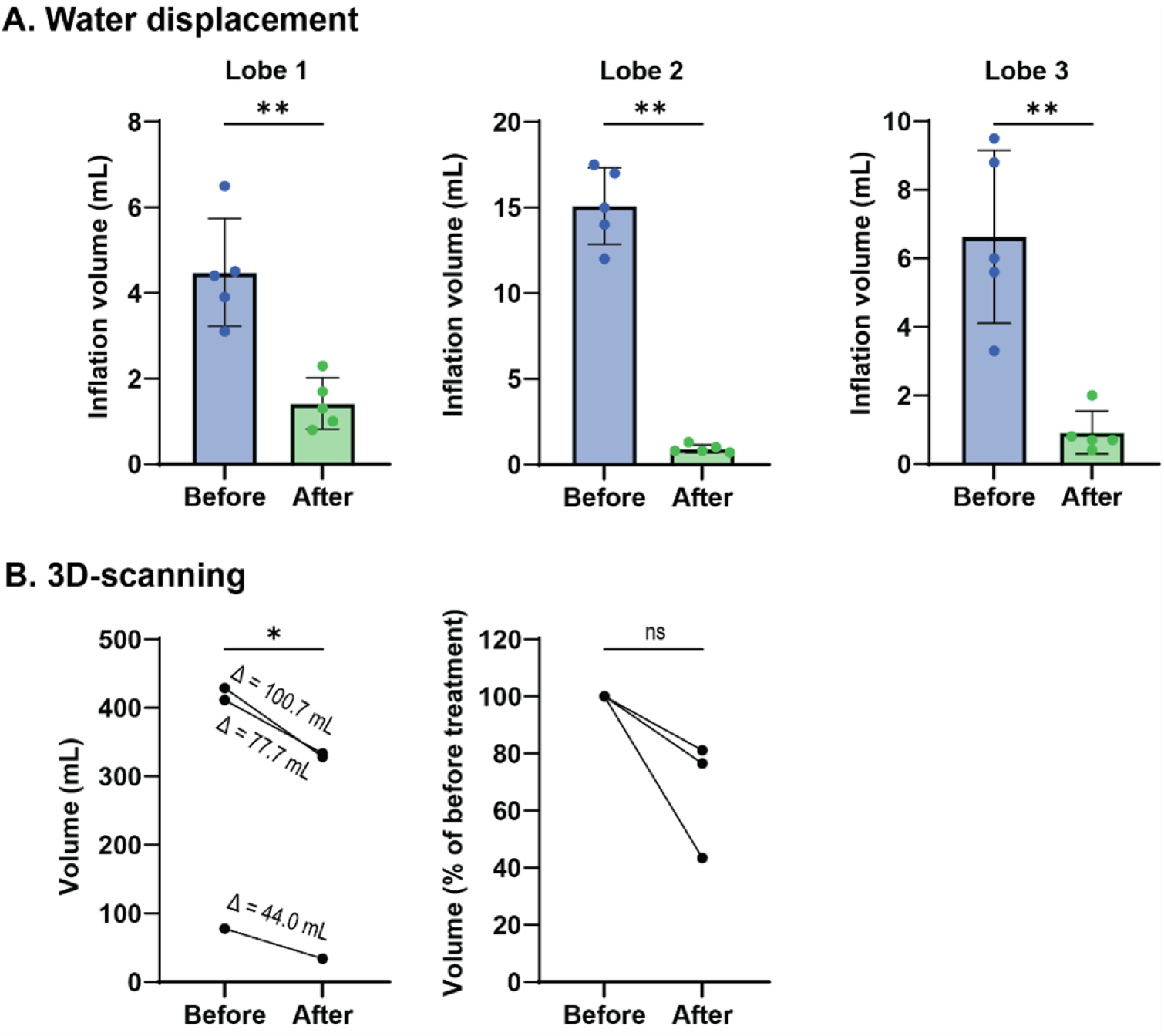
Results measured by water displacement and 3D scanning before and after treatment. (A) Water displacement method: volume increase upon inflation of three porcine lung lobes, before and after treatment with cyanoacrylate glue. Data shown as mean ± SD with n = 5 per condition, which are technical replicates (5 measurements before treatment and then 5 after treatment). Statistical differences were tested using a Mann–Whitney U test, **p ≤ 0.01. (B) 3D scanning method: volume of three porcine lung areas, before and after bronchoscopic administration of cyanoacrylate glue. Data shown in volumes with differences (Δ) noted in the graph (p=0,0459) and in percentages of before treatment (p=0,1088). Statistical difference were tested using a paired t-test.

To simulate a more physiologically relevant scenario, three whole lungs were treated bronchoscopically and scanned using a 3D scanner. The reconstructed volumes from the pre- and post-treatment target areas of the lung are shown in Figure 3B. The calculated volumes ranged between 77.84 to 429.14 mL for pre-treatment and 33.81 to 328.41 mL for post-treatment.

## Discussion

In this study, we have used water displacement and 3D scanning as methods to determine the volume reduction of *ex vivo* lung tissue. These techniques could be utilized in the future for early phase experiments for novel BLVR techniques, establishing an intermediate step between *in vitro* and *in vivo* testing. Translating from bench to clinical trial, this could be an appropriate model for addressing the performance of new BLVR agents before moving into *in vivo* animal models.

Employing the water displacement methodology provided consistent results for measuring *ex vivo* lung volume reduction. While 20mL of air was used to inflate the lobes, the inflation volume measured was lower, so not all of the air contributes to the increase in external volume. A possible cause can be low tissue compliance and leakage of air from the cut regions. The volume reduction after treatment is most likely due to obstruction of the airways, combined with hindered expansion of the parenchymal tissue. Overall, these physical changes in the lung tissues were reliably reflected in the water volume displaced when the lobes were inflated.

Using 3D scanning, we were also able to register lung volume reduction in a more clinically relevant scenario. The large variability of the results may be reduced using this method, while also complicating the setup.[9] The variation in the lobe volumes is apparent due to the biological heterogeneity in the porcine lung (difference in whole lung and lobe sizes). While minimal, there may also be some user-dependent variation in the measurements and post-processing of the 3D scans.[13] Lastly, despite our standardization, there could have been minimal differences in inflation status of the lung, which could influence volume measurements. Despite the variation, a consistent decline in volume after treatment was observed for each scanned area. This technique is therefore especially useful in measuring volume differences.

It is important to note that while both methods were able to observe a decrease in lung tissue volume, they are difficult to compare with each other head-on. Water displacement was performed using individual, separated lobes, while 3D scanning was performed using intact lungs. Both approaches are easy to apply and cost-effective. It must be taken into account that there exists inherent heterogeneity in the shape and size of the lungs and separated lobes, leading to variation in the results. Furthermore, for the 3D scanning method, it is crucial to perform the pre- and post-treatment scans at the same position and level of inflation; otherwise, alignment during analysis is not precise ultimately leading to less accurate measurements. Importantly, cyanoacrylate glue is not suitable for clinical BLVR due to its cytotoxicity, but was only utilized in order to demonstrate the validity of the quantification methods, ensuring robust differences before and after treatment by applying a well-recognized and effective adhesive.

In conclusion, this study showed that the water displacement and 3D scan methods reveal opportunities to objectively quantify lung volume reduction in *ex vivo* models, which is a large improvement compared to visual assessment. Using these methods, we aim to support the development of novel BLVR strategies while reducing the use of *in vivo* animal models.

## Acknowledgements

The authors thank Tim Abee for his assistance with the illustrations.

## Author contributions

All authors contributed to the study conception and design. Material preparation, data collection, and analysis were performed by ANK, SLSY, JAS, and DJS. The first draft of the manuscript was prepared by ANK and SLSY, and all authors commented on and edited versions of the manuscript. All authors read and approved the final manuscript.

## Data availability statement

All data are available from the authors upon request.

## Ethics statement

No human subjects or test animals were used for this study. All animal material used in this study is waste material derived from a commercial slaughterhouse.

## References

1. Snyder, L. E., Gonzalez, X., Barry, R. L., Pedersen, K. M. & Mulligan, M. S. Ex Vivo Human Lungs With Emphysema Used to Test a New Surgical System for Lung Volume ReductionFREE TO VIEW. Chest 124, 76S (2003).

2. Jiao, Y. et al. A preclinical animal study to evaluate the operability and safety of domestic one-way endobronchial valves. Front Med (Lausanne) 11, 1293940 (2024).

3. Ingenito, E. P. et al. Bronchoscopic Lung Volume Reduction Using Tissue Engineering Principles. Am J Respir Crit Care Med 167, 771–778 (2003).

4. Koster, T. D., Dijk, M. V. & Slebos, D.-J. ronchoscopic Lung Volume Reduction for Emphysema: Review and Update. Semin Respir Crit Care Med 43, 541–551 (2022).

5. Klooster, K. et al. Endobronchial Valves for Emphysema without Interlobar Collateral Ventilation. N Engl J Med 373, 2325–2335 (2015).

6. Come, C. E. et al. A randomised trial of lung sealant versus medical therapy for advanced emphysema. Eur Respir J 46, 651–662 (2015).

7. Niehues, S. et al. Liver volume measurement: reason of the difference between in vivo CT-volumetry and intraoperative ex vivo determination and how to cope it. Eur J Med Res 15, 345–350 (2010).

8. Lagendijk, M. et al. Breast and Tumour Volume Measurements in Breast Cancer Patients Using 3-D Automated Breast Volume Scanner Images. World J Surg 42, 2087–2093 (2018).

9. Haniuda, M. et al. Effects of inflation volume during lung preservation on pulmonary capillary permeability. The Journal of Thoracic and Cardiovascular Surgery 112, 85–93 (1996).

10. Chromy, A. High-Accuracy Volumetric Measurements of Soft Tissues using Robotic 3D Scanner. IFAC-PapersOnLine 48, 318–323 (2015).

11. Oezel, L. et al. Volumetry of Hand and Forearm: A 3D Volumetric Approach. Hand (N Y) 19, 845–850 (2024).

12. Schiltz, D. et al. Digital Volumetric Measurements Based on 3D Scans of the Lower Limb: A Valid and Reproducible Method for Evaluation in Lymphedema Therapy. Ann Vasc Surg 105, 209–217 (2024).

13. Schipper, J. a. M. et al. Reliability and validity of handheld structured light scanners and a static stereophotogrammetry system in facial three-dimensional surface imaging. Sci Rep 14, 8172 (2024).

14. Rush, S. A., Maddox, T., Fisk, A. T., Woodrey, M. S. & Cooper, R. J. A precise water displacement method for estimating egg volume. Journal of Field Ornithology 80, 193–197 (2009).

